# Chromosome-level assembly of *Cucumis sativus cv*. ‘Tokiwa’ as a reference genome of Japanese cucumber

**DOI:** 10.1101/2024.04.15.589484

**Authors:** Takashi Seiko, Chiaki Muto, Koichiro Shimomura, Ryoichi Yano, Yoichi Kawazu, Mitsuhiro Sugiyama, Kenji Kato, Norihiko Tomooka, Ken Naito

## Abstract

Cucumber is one of the most important vegetables in the Japanese market. To facilitate genomics-based breeding, there is a demand for reference genome of Japanese cucumber. However, although cucumber genome is relatively small, its assembly is a challenging issue because of tandem repeats comprising ∼30% (∼100 Mbp) of the genome. To overcome, we deployed the Oxford nanopore sequencing that produces long reads with N50 length of >30 kbp. With this technology we achieved a chromosome-level assembly of cv ‘Tokiwa’, a founder line of Japanese cucumber represented with the elongated fruit shape and high-crisp texture. Compared to the existing cucumber genomes, the Tokiwa genome is 20% longer and annotated with 10% more genes. The assembly with nanopore long reads also resolved tandem repeats spanning >100 kbp, demonstrating its strength in overcoming repetitive sequences.

## Introduction

Cucumber (*Cucumis sativus* L.) shares the 3rd place in the value production in the Japanese vegetable market, after tomato and onion (https://www.usdajapan.org/wpusda/wp-content/uploads/2019/03/Japanese-Fresh-Vegetable-Market-Overview-2018_Osaka-ATO_Japan_12-21-2018-1.pdf). The Japanese cultivars of cucumber are represented with a long and straight fruit shape (Shimomura et al. 2016) and a crispy texture (Yoshioka et al., 2009), due to specific needs of the customers. Thus, breeding efforts of cucumber mainly target resistance to diseases including melon yellow spot virus and powdery mildews without spoiling the important traits described above. Heat tolerance has also become an important target because of the global warming. To facilitate breeding of such cultivars, we need to reconstruct a reference genome of Japanese cucumber and implement marker-assisted selection, allele mining *via* GWAS and prediction models for genomic selection.

However, efforts to sequence cucumber genomes have faced difficulties despite its relatively small size (∼350 Mbp) (Huang et al., 2009, Yang et al., 2012, Li et al., 2019). The current reference genome is “Chinese long” inbred line 9930, which was sequenced with single molecule real-time (SMRT) sequencing technology (Li et al., 2019). The 9930 genome consists of 7 pseudomolecules covering 211 Mbp + 15 Mbp unanchored scaffolds, leaving ∼120 Mbp sequences missing (Huang et al., 2009). More recently 11 more accessions have also been sequenced with the SMRT sequencing, but the assemblies were only slightly better than the 9930 genome (Li et al., 2022). The difficulty in assembling cucumber genome derives from long arrays of tandemly repeated sequences around centromeric regions (Ganal et al., 1986, Ganal and Hemleben, 1988).

To overcome the difficulties, increasing read length is the simplest solution. We adopted a recently-developed technology by Oxford Nanopore, which can generate ultra long reads spanning more than 100 kbp (Wang et al, 2021). Many plant genomes have already been sequenced with this technology, some achieving the NG50 length of more than 10 Mbp (Dumschott et al., 2020). Thus, we expect it may improve *de novo* assembly of a cucumber genome.

Here, we sequenced and assembled the whole genome of a Japanese cucumber cultivar, cv. ‘Tokiwa’ (Fig. 1). ‘Tokiwa’ was developed from a cross between North China type ‘Shimo-shirazu’ and South China type ‘Shindome’ around 1960. Given its Japanese favored fruit morphology and texture and adaptability to both summer and fall cropping system, it has been a founder line of various modern cultivars, which is now called as “Tokiwa group” (Fujieda, 2006). The Nanopore-only assembly generated greatly improved contiguity and coverage compared to the 9930 genome, providing a qualified knowledge-base for Japanese breeders.

**Figure 1.**
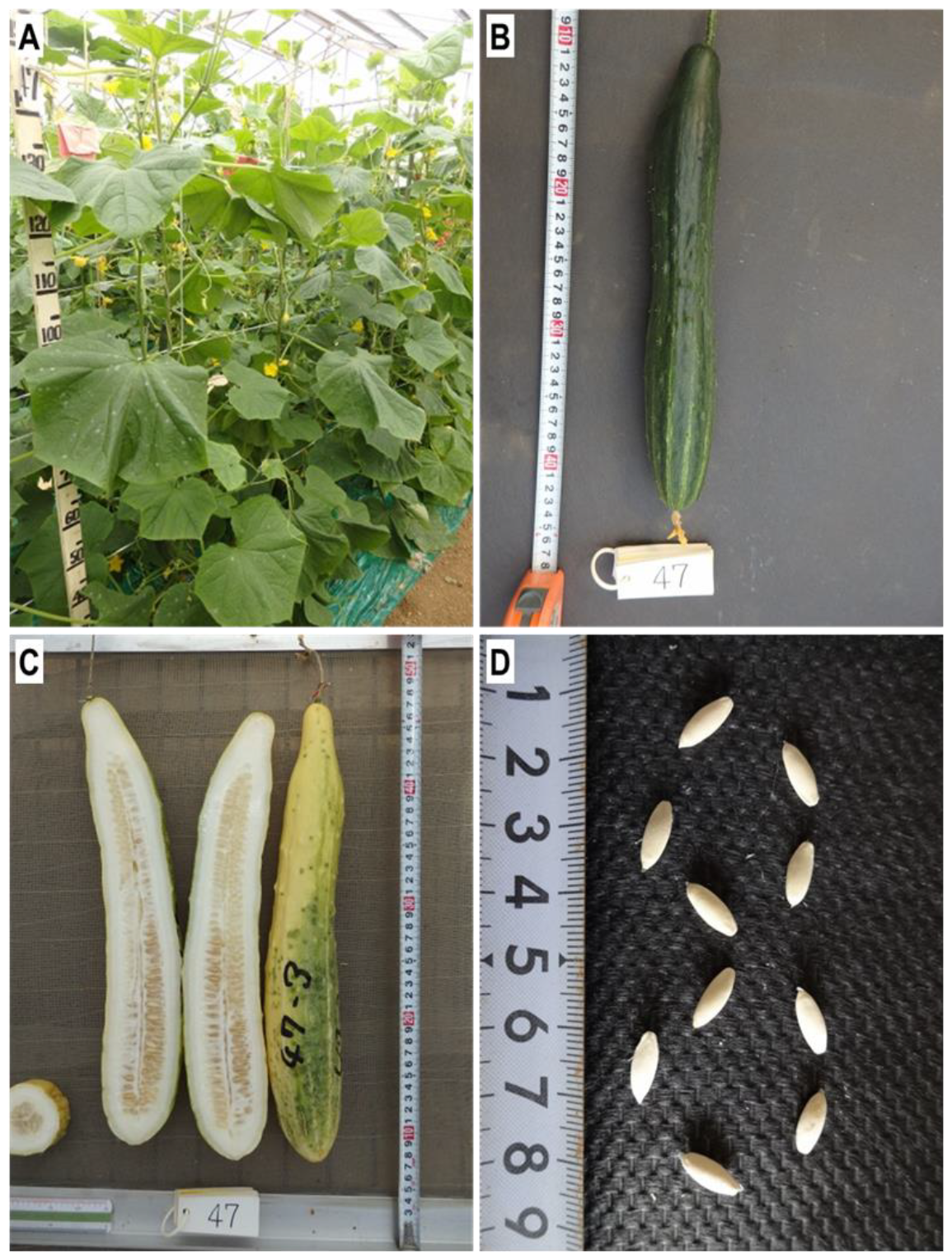
Photographs of *Cucumis sativus* L. cv ‘Tokiwa’. a. Plants growing in a greenhouse, b. A green fresh fruit, c. Mature fruites. d. Seeds.

## Materials and methods

### Plant material and DNA extraction

The cucumber cultivar ‘Tokiwa’ was provided by National Genebank Program in Research Center of Genetic Resources, NARO. We extracted genomic DNA from 3 g of young leaves using Nucleobond HMW DNA kit (MACHEREY-NAGEL GmbH & Co. KG, Düren, Germany). We basically followed the DNA extraction protocol provided by the manufacturer, but we extended the lysis step up to 3 h instead of 30 min. In the precipitation + washing step with isopropanol and 70% (v/v) ethanol, we did not centrifuge but directly transfer the precipitated DNA by pipetting.

We also extracted RNA from the leaves for gene annotation used by NucleoSpin RNA Plant kit (MACHEREY-NAGEL GmbH & Co. KG, Düren, Germany).

### Sequence of DNA and RNA

For long-read sequencing, we followed the protocol as described by Naito (2023). First, we size-selected the genomic DNA using Short Read Eliminatior XL kit (Pacific Biosciences of California, Inc., California,USA) to remove the DNA fragments <10 kbp. Second, we put 3 μg of the size-selected DNA for end-repair using NEBNext FFPE DNA Repair Mix (NEW INGLAND BioLabs inc., Massachusetts, USA) and NEBNext Ultra II End Repair/dA-Tailing Module. Third, we ligated the end-repaired DNA with the sequencing adaptor in SQK-LSK109 (Oxford Nanopore Technologies KK, Tokyo, Japan) using NEBNext Quick Ligation Module (NEW INGLAND BioLabs inc., Massachusetts, USA). In each step, we purified DNA using AMPure XP (BECKMAN COULTER inc., California, USA). The prepared library was then loaded to R9.4.1 Flowcell (Oxford Nanopore Technologies KK, Tokyo, Japan) and was run in PromethION24 (Oxford Nanopore Technologies PLC, Oxford, UK) for 10 h. We also obtained short-reads for polishing the draft assembly with Illumina Hiseq 4000, which was provided as a customer service by GeneBay, Inc. (Yokohama, Japan). We also sequenced the RNA with Hiseq4000 (Illumina Inc., San Diego, USA) for gene annotation, provided as a customer service by GeneBay Inc. (Yokohama, Japan). All the data is uploaded to the Sequence Read Archive in NCBI website, with BioSample ID SAMN40100976 in BioProject ID PRJNA1080095.

### Genome assembly and scaffolding

We also followed the protocol described by Naito (2023) for genome assembly. Fist, we used necat-0.0.1 (Chen et al., 2021) with default parameters except “PREP_OUTPUT_COVERAGE=60” and “CNS_OUTPUT_COVERAGE=40”. The draft contigs were then polished twice with racon-1.4.3 (Vaser et al., 2017) with options of “-m 8 -x -6 -g -8 -w 500” and medaka-1.0.3 (https://github.com/nanoporetech/medaka) with options of “-g -m r941_prom_high_g360”. To check misassemblies, we first aligned the polished contigs to the 9930 genome by D-genies (Cabanettes and Klopp 2018) to visualize conflicted sites. We then mapped the long-reads with minimap2-2.14 (Li, 2018) and manually checked the conflicted sites. If the mapped reads were discontinuous as presented by Sakai et al. (2015), we broke the contig at the site. The corrected contigs were further polished with Illumina short-reads twice with Hypo-1.0.3 (https://github.com/kensung-lab/hypo) with default parameters. The Hypo-polished contigs were then run with purge_haplotigs-1.1.1 (Roach et al., 2018) to remove low-coverage contigs and those derived from heterozygous regions.

After assembling, polishing and filtering the contigs, we obtained optical mapping data provided as a customer service by AS ONE, corp. (Osaka, Japan). High-molecule genomic DNA was extracted from young leaves with the Plant DNA Isolation Kit (Bionano Genomics, San Diego, CA, USA) according to Bionano Prep Plant Tissue DNA Isolation Base Protocol. The DNA was treated with DLE-1 nickase, labelled with DLS DNA Labeling Kit (Bionano Genomics), and run on the Saphyr Genome Mapping Instrument (Bionano Genomics). The obtained Saphyr reads were assembled and then merged with the draft assembly to generate hybrid scaffolds sequences with Bionano Solve (Bionano Genomics) with the default parameters. Tha Bionano data is also available from NCBI website with BioProject ID: PRJNA1098469.

Finally, pseudomolecules were reconstructed by anchoring the hybrid scaffolds to the genetic map, which was previously constructed (Sugiyama et al., 2016).

To compare the assembly with 9930 genome, we made a whole genome alignment by minimap2 and ran the output with D-genies to generate a dotplot.

### Assembling organelle genomes

Given there are hundreds of chloroplasts and tens of mitochondria in a single cell, the coverage depth is usually too high for assembling the organellar genomes. Thus, we had to extract and downsample the long reads derived from the organellar genomes.

To do so, we first identified the organellar contigs by BLASTing the reference genomes of chloroplast (Pląder et al., 2007) and mitochondrion (Alverson et al., 2011) as query to the draft assembly. Then we mapped the long reads to the draft assembly and recovered the reads that were mapped to the organellar contigs. We then downsampled the recovered reads to ∼50x coverage and ran Flye-2.9 (Kolmogorov et al., 2019). The assembled sequences were then polished with long reads with medaka.

### Gene predictions

Briefly, we predicted gene structures using Braker-2.0 (Brůna et al., 2021) using protein sequences and GeMoMa-1.9 (Keilwagen et al., 2018) using our own RNA-seq data and the coding sequences of 9930 genome.

First, we softmasked the Tokiwa genome by running RepeatModeler-2.0.5 (https://www.repeatmasker.org/RepeatModeler/) and RepeatMasker-4.0.9 (https://www.repeatmasker.org/RepeatMasker/) with default parameters. Second, we downloaded the sequences of plant proteins from OrthoDB (Kuznetsov et al., 2022) as well as the license for GeneMark-EP+ (Brůna et al., 2020), which was required for running Braker-2.0. Third, we ran braker-2.0 on the softmasked genome with an option of “--softmasking”.

Before running GeMoMa, we removed noisy reads from the RNA-seq data. To do so, we *de novo* assembled the RNA-seq data with Trinity-2.13.0 (Grabherr et al., 2011) with default parameters. Then we mapped the RNA-seq reads to the assembled transcript sequences using HISAT-2.2.1 (Kim et al., 2019) and recovered only concordantly mapped reads. The recovered RNA-seq reads were then mapped to Tokiwa genome by HISAT-2.2.1 with an option of “--max-intronlen 15000”, which was used for intron prediction by GeMoMa. Then we ran GeMoMa using 9930 genome and gene annotations as reference, with parameters for unstranded RNA-seq data. After running the annotation tools, we merged the outputs of GeMoMa and Braker, by filtering the overlapping annotations.

Then we ran EnTAP-0.10.8 (Hart et al., 2020) with default parameters to functionally annotate the predicted genes. The final annotations retained those identified as “Viridiplantae” and “Eukaryotes” for Taxonomy scope.

We also reannotated the 9930 genome and those of the 11 accessions assembled by Li et al. (2022). The 11 genome sequences were downloaded from the website of NCBI datasets. The downloaded genomes were annotated by GeMoMa-1.9 with the predicted gene models of ‘Tokiwa’ as reference. JCVI MCscan (https://github.com/tanghaibao/jcvi/wiki/MCscan-(Python-version)) was used to predict synteny block between each genome.

The predicted genes were evaluated with eudicots_odb10 (Kriventseva et al., 2018) by BUSCO-5.5 (Manni et al., 2021).

### Phylogenetic analysis

To place the Tokiwa genome in a phylogenetic position, we first ran orthofinder-2.5.5 (Emms et al., 2019) on the protein sequences of the 13 accessions, with “-M msa” option. Then we reconstructed a maximum likelihood tree from the ouput mafft file of single copy orthologs by running iqtree-2.2.6 with options of “-m MFP -bb 1000 -alrt 1000 -nt AUTO”.

### Repeat annotation

To identify sequences derived from transposable elements, we ran EDTA-2.2.2 (Ou et al., 2019) with the predicted coding sequences of ‘Tokiwa’ in a sensitive mode. To identify tandem repeats, we ran trf-4.0.9 with parameters suggested by the official protocol.

## Results

### De novo *Assembly of Tokiwa genome*

We have successfully reconstructed the pseudomolecules of the 7 chromosomes in Tokiwa genome, covering 50 Mbp more than the 9930 genome (Table 1, Fig. 2).

**Table 1.** The statistics of genome assemblies.

|  | NECAT | Purge_haplotigs | BioNano | Tokiwa v1.0 | 9930 v3 |
| --- | --- | --- | --- | --- | --- |
| Num | 645 | 385 | 314 | 7 | 7 |
| Sum | 300,877,148 | 278,732,282 | 299,675,636 | 257,281,677 | 210,974,704 |
| Min | 208 | 4,095 | 4,095 | 27,903,525 | 22,466,726 |
| Ave | 466,476 | 723,980 | 954,381 | 36,754,525 | 30,139,243 |
| Max | 20,967,667 | 20,967,667 | 45,117,507 | 45,117,507 | 40,877,379 |
| N50 | 12,103,241 | 12,256,096 | 23,376,839 | 36,771,724 | 31,913,682 |
| GAP | - | - | 20,923,098 | 15,419,875 | 36,170 |
| Total ATGC | - | - | 278,752,538 | 241,861,802 | 210,938,534 |
| Non-placed scaffolds | - | - | - | 130 | 81 |
| Non-placed bps | - | - | - | 23,587,426 | 15,666,262 |
| Protein coding genes |  |  |  | 25,159 | 23,660 |

**Figure 2.**
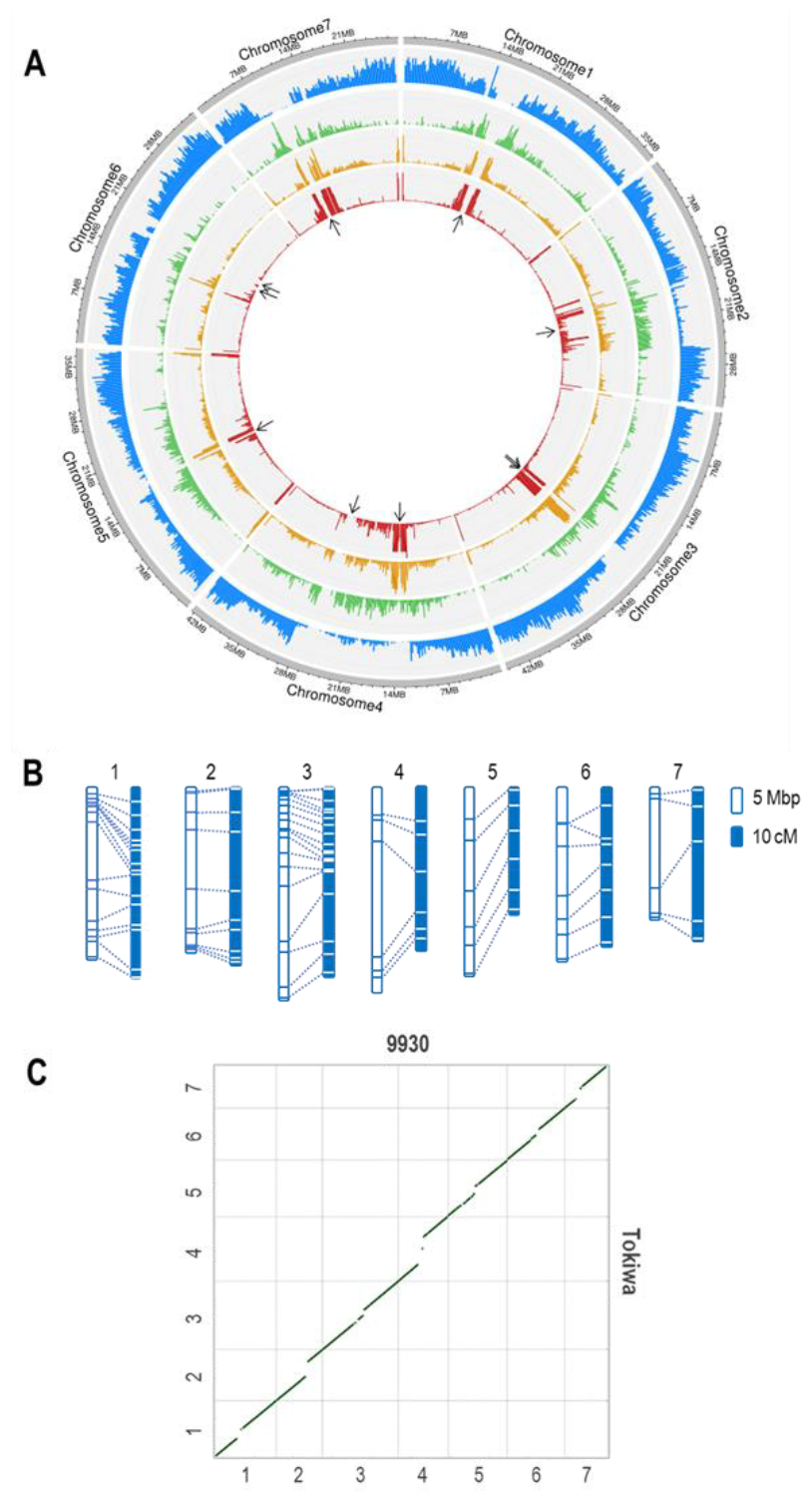
Overview of Taokiwa genome. A. Circos plot indicating densities of, from the outer to inner tracks, protein coding sequences, LTR retrotransposons, DNA transposable elements, and tandem repeats. B. Coliniarity between the physical positions and the genetic linkage map of the Tokiwa genome. White and blue boxes indicate the physical and genetic map, respectively. Dashed lines indicate the corresponding positions. C. Dotplot of a whole genome alignment of 9930 and Tokiwa. *x*- and *y* -axes indicate Chromosome numbers. Cultivar names are indicated above the x-axis and right to the *y*-axis.

Using the Oxford Nanopore sequencer, we obtained 34.2 Gbp of long reads with the N50 length of 33.2 kbp (Supplemental Table 1). The draft assembly with NECAT was ∼300 Mbp sequences in 645 contigs, with the N50 length of 12.1 Mbp and the maximum length of 20.1 Mbp (Table 1). Purge_haplotigs identified 260 contigs as repetitive or low-coverage junk and thus retained 385 contigs covering 278.7 Mbp. The following optical mapping scaffolded only 71 contigs but improved the N50 length to 23.4 Mbp and the maximum length to 45.1 Mbp. Finally, we anchored the 14 scaffolds to the genetic linkage map (Sugiyama et al., 2015) and reconstructed the pseudomolecules of the 7 chromosomes (Fig. 2). After removing the scaffolds/contigs derived from organelle genome and possible contaminations, 23.6 Mbp in 130 scaffolds/contigs remained unplaced. All the seven pseudomolecules contained telomeric repeats in both ends.

We also separately assembled the genome sequences of the chloroplast and mitochondrion. The chloroplast genome was successfully assembled into a single circular genome. However, due to the highly recombining feature of mitochondrion, we could not circularize the mitochondrial genome and ended up with 6 contigs (Supplemental Table 2).

### More duplicated genes predicted in the Tokiwa genome

The pseudomolecules of Tokiwa genome reached 257 Mbp, which was >40 Mbp longer than those of the 9930 genome (Fig. 2B). Even though the extended sequences were mostly in centromeric, highly repetitive regions, gene coding sequences were also present (Fig. 2A). Although we used the Tokiwa’s gene models to reannotate the 9930 and other genomes for fair comparison, the Tokiwa genome contained ∼2,000 more genes than others (Table 2). The BUSCO scores were not greatly different among them, but the predicted gene models in the Tokiwa genome contained the fewest fragmented or missing BUSCO genes (Table 2).

**Table 2.** BUSCO scores of the Tokiwa other genomes.

|  | Complete |  | Single |  | Duplicate |  | Fragmented |  | Missing |  |
| --- | --- | --- | --- | --- | --- | --- | --- | --- | --- | --- |
|  | Number | (%) | Number | (%) | Number | (%) | Number | (%) | Number | (%) |
| Tokiwa | 2,275 | (97.8) | 2,217 | (96.0) | 43 | (1.8) | 10 | (0.4) | 41 | (1.8) |
| 9930 | 2,268 | (97.5) | 2,217 | (95.3) | 51 | (2.2) | 12 | (0.5) | 46 | (2.0) |
| 9110gt | 2,250 | (96.7) | 2,213 | (95.1) | 37 | (1.6) | 11 | (0.5) | 65 | (2.8) |
| Cu2 | 2,266 | (97.4) | 2,212 | (95.1) | 54 | (2.3) | 10 | (0.4) | 50 | (2.2) |
| Cuc37 | 2,262 | (97.3) | 2,221 | (95.5) | 41 | (1.8) | 11 | (0.5) | 53 | (2.2) |
| Cuc64 | 2,273 | (97.7) | 2,231 | (95.9) | 42 | (1.8) | 8 | (0.9) | 45 | (2.0) |
| Cuc80 | 2,272 | (97.7) | 2,225 | (95.7) | 47 | (2.0) | 8 | (0.3) | 46 | (2.0) |
| Gy14 | 2,256 | (97.0) | 2,212 | (95.1) | 44 | (1.9) | 10 | (0.4) | 60 | (2.6) |
| Hx14 | 2,251 | (96.7) | 2,213 | (95.1) | 38 | (1.6) | 11 | (0.5) | 64 | (2.8) |
| Hx117 | 2,233 | (96.0) | 2,189 | (94.1) | 44 | (1.9) | 15 | (0.6) | 78 | (3.4) |
| W4 | 2,267 | (97.4) | 2,224 | (95.6) | 43 | (1.8) | 9 | (0.4) | 50 | (2.2) |
| W8 | 2,251 | (96.7) | 2,206 | (94.8) | 45 | (1.9) | 13 | (0.6) | 62 | (2.7) |
| XMTC | 2,259 | (97.1) | 2,217 | (95.3) | 42 | (1.8) | 12 | (0.5) | 55 | (2.4) |

The genome annotation revealed highly conserved synteny between the cucumber genomes including wild accessions, except several inversions in Chromosomes 4, 5, and 7 of Cuc64 and W8, which were also revealed by genome alignment-based analysis (Li et al., 2022). (Fig. 3). However, it also revealed that the genes present in the newly assembled regions of the Tokiwa genomes were mostly the duplicated copies of those in other regions (Fig. 3).

**Figure 3.**
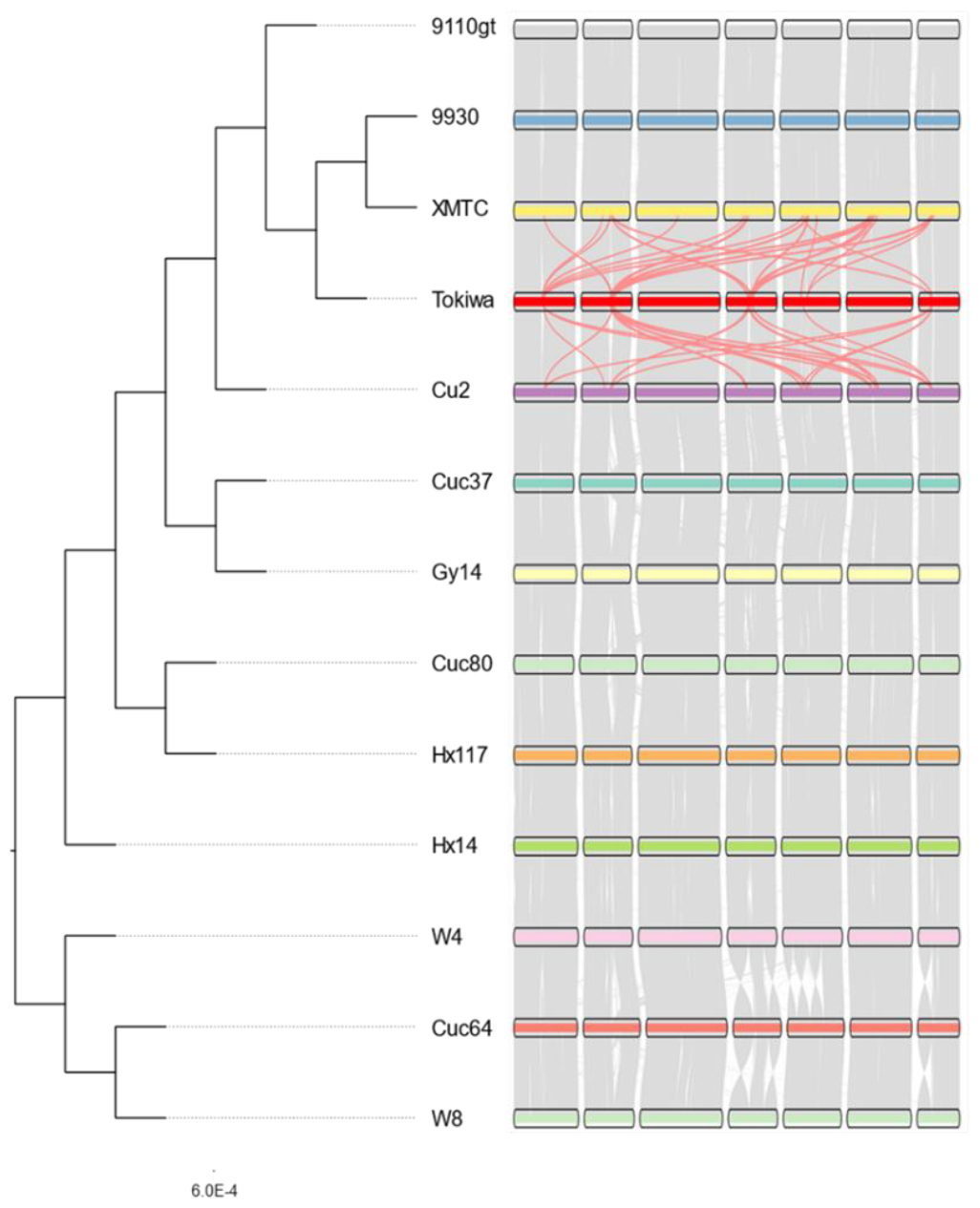
Phylogenetic tree and synteny plot of the 12 chromosome-level genome assemblies. Gray lines indicate synteny blocks between the genomes. Light red lines indicate pseudo synteny blocks between the newly assembled regions of the Tokiwa genome and XMTC or Cu2. Bootstrap values were 100 in all the nodes.

The following ortholog analysis enabled us to predict the number of duplicated genes. Although more than 90% of the predicted genes were present as single copy in the 13 accessions, the Tokiwa genome had the highest number of genes (much higher number of those present as multi-copy orthologs (Supplementary Table 3).

The ortholog analysis also enabled us to extract 17,123 single-copy orthologs and reconstruct a phylogenetic tree of the 13 accessions (Fig. 3). It grouped the cultivars (9110gt, 9930, XMTC, Tokiwa, Cu2, Cuc37, Gy14, Cuc80, Hx117 and HX14) and the wild cucumbers (*C. sativus* var. *hardwickii*) (W4, Cuc64 and W8) into monophyletic clades, respectively, placing ‘Tokiwa’ in the East-Asian group together with the Chinese cultivars 9110gt, 9930, XMTC, and Cu2 (Fig. 3).

### Distribution of transposable elements in Tokiwa genome

Because of the relatively small size of the genome, transposable elements shared only 23.9% of the Tokiwa genome (Supplemental Table 4). Overall, DNA transposons were more abundant than LTR retrotransposons were. The most abundant was Mutator family, comprising nearly 10% (22.8 Mbp) of the Tokiwa genome, followed by Gypsy, Copia, and CACTA.

In the Tokiwa genome, the distribution of LTR retrotransposons and DNA transposons were basically exclusive to each other (Fig. 2A). While the DNA transposons were highly condensed near centromeric and the telomeric regions, the LTR retrotransposons were more broadly distributed across the pericentromeric regions.

### Tandem repeat in Tokiwa genome

As cucumber genome is peculiar for harboring long stretches of tandem repeats, we also surveyed the Tokiwa genome with tandem repeat finder (Benson 1999). The survey revealed that tandem repeats comprise 35.2 Mbp (12.6%) of the Tokiwa genome.

Although the size of the repeat unit varied, those of ∼180 bp, ∼360 bp, ∼540 bp and ∼720 bp had extremely high copy numbers as well as those <10 bp (Fig. 4). The longest stretch of the tandem repeats was that of 712.6 kbp on Chromosome 4, where a unit of 531 bp sequence was tandemly repeated by 1,331 times (Table 3). Other chromosomes also harbored tandem repeats spanning >100 kbp (Table. 3), which could be even longer, as many of them ended with gaps.

**Figure 4.**
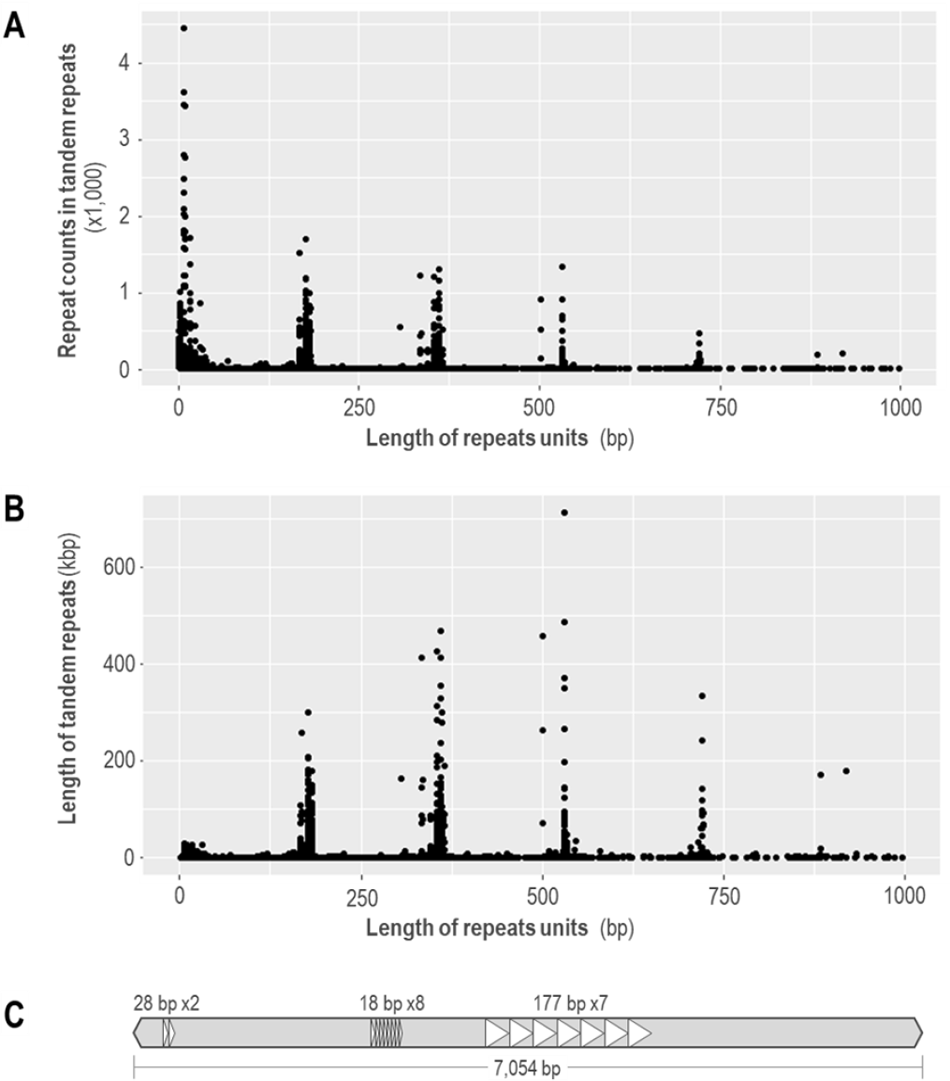
Distribution of unit size, repeat count and total length of the tandem repeats in the Tokiwa genome. A. Scatter plot of unit size (*x*-axis) and repeat count (*y*-axis) of each tandem repeat. B. Scatter plot of unit size (*x*-axis) and the total length of each tandem repeat (*y*-axis). C. Schematic of a Mutator-like element containing tandem repeats. Triangles indicate the units of each tandem repeat.

**Table 3.** The top 10 longest tandem repeats in the Tokiwa Genome.

| Chromosome | Start | End | Unit<br>size | Repeat<br>Count | Length |
| --- | --- | --- | --- | --- | --- |
| Chromosome4 | 12,049,307 | 12,761,914 | 531 | 1331.1 | 712,608 |
| Chromosome3 | 25,304,414 | 25,792,696 | 531 | 915.6 | 488,283 |
| Chromosome5 | 109,973 | 578,952 | 360 | 1301 | 468,980 |
| Chromosome1 | 14,073,064 | 14,531,408 | 501 | 903.2 | 458,345 |
| Chromosome7 | 7,938,677 | 8,366,883 | 354 | 1203.4 | 428,207 |
| Chromosome7 | 8,006,713 | 8,377,207 | 531 | 693.4 | 370,495 |
| Chromosome7 | 27,538,573 | 27,895,514 | 360 | 989.3 | 356,942 |
| Chromosome7 | 10,064,253 | 10,414,004 | 531 | 654.1 | 349,752 |
| Chromosome5 | 37,052,513 | 37,392,146 | 720 | 463.2 | 334,450 |

Interestingly, the long stretches of tandem repeats (>10 kbp) were often overlapped with annotations of Mutator and CACTA transposons (Fig 2). In addition, the Mutator and CACTA elements identified by EDTA often contained the tandem repeats as their internal sequences (Fig. 4C). Namely, the tandem repeats did not cover the full length of the transposable elements. Thus, increase in tandem repeats did not necessarily occur *via* TE amplifications.

## Discussion

Here we present the most contiguous assembly of the cucumber genome, owing to the long reads by Oxford Nanopore. The assembled Tokiwa genome is 50 Mbp longer and predicted 2,000 more gene coding sequences compared to the 9930 genome or other existing assemblies (Table 2, Figs. 2, 3, Supplementary Table 3). The annotation predicted more duplicated genes, less fragmented genes, and much more repeat sequences including tandem repeats. As ‘Tokiwa’ is a founder line of modern Japanese cultivars, the highly contiguous and accurate genome could facilitate breeding for urgent and long-term demands in Japanese market.

Our results have demonstrated the importance of read length to overcome the repetitive sequences in *de novo* assembly. The nanopore sequencer produced long reads with N50 length of 33.2 kbp, which was >3 time longer than those used for assembling the 9930 genome (Supplementary Table 1). Such long reads have enabled us to resolve and assemble long stretches of tandem repeats, including the one over 700 kbp (Table 3).

However, we should also note that we could not assemble centromeric regions without the optical mapping by Bionano. Although the draft assembly already generated contigs containing long stretches of tandem repeats, such contigs could not be directly anchored to the linkage map because there were no genetic markers in such highly repetitive regions. Thus, optical mapping was the only way to place the repeat-rich contigs in centromeric or pericentromeric regions.

The newly resolved repeat sequences also give insight into the relationship between the DNA transposons and the tandem repeats. The distribution of the unit size in the tandem repeats indicated that those with high copy numbers are overrepresented with specific unit sizes. This led us to suspect that such tandem repeats could arise from cis-transposition of transposable elements. However, many of the tandem repeats are actually “internal” sequences of transposons. Namely, one single copy of a transposon could be very long, containing a long stretch of tandem repeats within its sequence (Fig. 4). Thus, the tandem repeats could have increased its copy number via other mechanisms than transposition, such as illegitimate recombination or double strand break repair.

Although the long reads by Oxford Nanopore greatly facilitated assembling highly repetitive regions, the Tokiwa genome is still far from gapless assembly. Given the estimated size of cucumber genome is ∼350 Mbp, there remains ∼70 Mbp sequences unassembled. As the Oxford Nanopore have greatly improved the accuracy of raw reads (Zhao et al., 2023), new assemblers optimized for lower error rate would further facilitate resolving the tandem repeats.

To conclude, this study presented a *de novo* assembly of Tokiwa genome using long reads of Oxford Nanopore, which could serve as a reference of Japanese cucumber. We hope the Tokiwa genome would be a fundamental knowledge-base for future breeding of Japanese cultivars.

## Author’s contributions

**KN, NT** and **KK** conceived the study.

**TS** and **CM** performed sequencing and assembly.

**CM, KS, YK, MS, RY** and **KN** fixed misassemblies and performed scaffolding.

**TS** and **KN** performed genome annotation and data analysis.

**TS, KS** and **KN** wrote the paper.

All the authors read and approved the final manuscript.

## Data availability

All the sequence data were deposited at Biosample IDs SAMN40100976, under Bioproject PRJNA1080095.

